# Structural and functional MRI from a cross-sectional Southwest University Adult lifespan Dataset (SALD)

**DOI:** 10.1101/177279

**Authors:** Dongtao Wei, Kaixiang Zhuang, Qunlin Chen, Wenjing Yang, Wei Liu, Kangcheng Wang, Jiangzhou Sun, Jiang Qiu

## Abstract

Recently, the field of developmental neuroscience has aimed to uncover the developmental trajectory of the human brain and understand the changes that occur as a function of aging. Here we present an adult lifespan dataset of functional magnetic resonance imaging (fMRI) data including structural MRI and resting-state functional MRI. 494 healthy adults (age range: 19-80 years; Males=187) were recruited and completed two multi-modal MRI scan sessions in the Brain Imaging Central of Southwest University, Chongqing, China. The goals of the dataset are to give researchers the opportunity to map the developmental trajectory of structural and functional changes of human brain and to replicate previous findings.

## Background & Summary

Magnetic resonance imaging (MRI) has been one of the most dominant techniques to investigate the human brain, because it permits the detailed, noninvasive and safe assessment of human brain. MRI is also able to perform the data collection of various image modalities, such as structural magnetic resonance imaging (sMRI), functional MRI (fMRI) and diffusion tensor imaging (DTI). In particular, these imaging measurements have been effectively used to capture brain structural and functional changes in development^1^, aging^2^, psychiatric disorders^3^, etc. For example, the feature of resting-state functional connectivity can predict the maturity of the individual across development^4^, as well as can be used as a “fingerprint” to identify individuals^5^. Thus, the measurements of MRI have greatly contributed to be served as imaging biomarkers of normal development, aging, clinical diagnosis and therapeutic assessments.

One of the most urgent scientific issue confronting us in the 21^st^ century is how can we maintain a healthy mind for human life. Besides paying attention to uncover the developmental course an original brain grows up to a mature one, another critical question in lifespan developmental neuroscience is how the brain changes as a function of aging. There is a necessity to answer this question, because only if we reveal the healthy brain aging mechanism can we discover the causes of brain diseases relating to aging (e.g., Alzheimer’s disease). Based on measurements of various image modalities, researchers have uncovered many appealing findings in the normal aging brain. For example, most brain regions follow a liner decline of gray matter volume (GMV) with normal aging, while nonlinear age trajectories were also observed in some regions (e.g., medial temporal lobe), which indicated a preservation of GMV during the early adult lifespan^6^; increasing age was found accompany by decreasing functional segregation of brain systems, and this age-related effect was more prominent in associative systems than in sensory and motor systems^7^.

For the sake of characterizing age-related changes in cognition and brain structure and function, the data repository for the Cambridge Centre for Ageing and Neuroscience (Cam-CAN) initial study cohort has yet provided a multi-model dataset from a large, cross-sectional adult lifespan population-based sample^8^. In addition, there are a number of publicly available datasets for free to authorized investigators, such as Open Access Series of Imaging Studies (OASIS) and Alzheimer’s Disease Neuroimaging Initiative (ADNI, (http://adni.loni.usc.edu/data-samples/). OASIS consists of a cross-sectional collection of 416 subjects aged 18 to 96and a longitudinalMRI Data collection of 150 subjects aged 60 to 96 (http://www.oasis-brains.org/). However, there is still lack of the open access dataset expanding beyond Caucasian white population, and allowing researchers to discover meaningful regulations of normal brain aging or verify previous findings. Moreover, to reveal age-related changes of human brain should be based on large continuous samples, which in a way limit research activities in aging. Thus, an additional open access normal adult lifespan data with the large sample is needed for researchers who are interested in this domain or require an independent dataset for cross-validation. Here, we describe the data generated in the Southwest University Adult Lifespan dataset (SALD), which is one part of our ongoing project to examine the association among brain imaging, creativity and mental health (BCM). The SALD comprises a large cross-sectional sample (total scans = 494; age span = 19-80 years), multi-modal (sMRI and rs-fMRI) investigation of the neural underpinnings. The goal of the SALD is to understand what a normal brain looks like and how it structurally and functionally changes at each decade of life from age 20 through 80. Now, it is available for research through the International Data-sharing Initiative (INDI, http://fcon_1000.projects.nitrc.org/indi/retro/sald.html). We hope our free data sharing can speed the progress of normal brain aging studies.

## Methods

### Participants

The 494 participants (308 Females, 187 Males, aged 19 to 80) included in the release were selected from a large dataset of individuals who have participated in the ongoing BCM data collection initiative. The young adults (18-25) of the lifespan sample enrolled as college students of Southwest University in Chongqing, China. Southwest University is a key comprehensive university affiliated to the Ministry of Education, by means of the merge of the former Southwest Normal University and Agricultural University of Southwest. The university enrolls 10, 000 ordinary undergraduates each year, nearly 65% are female. The young adultswere collected by random sampling from Southwest University, thus, the number of the young and female (20-27)is larger in our dataset(for more details, see figure 1). Many of the mid adults (age 26 to 40) were recruited directly from staff of Southwest University. The rest of adults sample were recruited from communities close to the university campus. The data collection was initiated in 2010 and was terminated in 2015. In addition, a part of participants serves as a control sample in a case-control study of a clinical population. Some participants were excluded due to sleeping during scanning. We primarily recruited participants through leaflets, online advertisements, and face-to-face propaganda. The exclusion criteria included: (1) MRI related exclusion criteria, which included claustrophobia, metallic implants, Meniere’s Syndrome and a history of fainting within the previous 6 months; (2) current psychiatric disorders and neurological disorders; (3) use of psychiatric drugs within the three months prior to scanning; (4) pregnancy; or (5) a history of head trauma. Informed written consent was obtained from each participant. Besides, we required participants to refrain from drinking during the day before the scanning and the scanning day. The dataset collection was approved by the Research Ethics Committee of the Brain Imaging Center of Southwest University in accordance with the Declaration of Helsinki. Written informed consent was obtained from all participants prior to the data collection.

**Figure 1.**
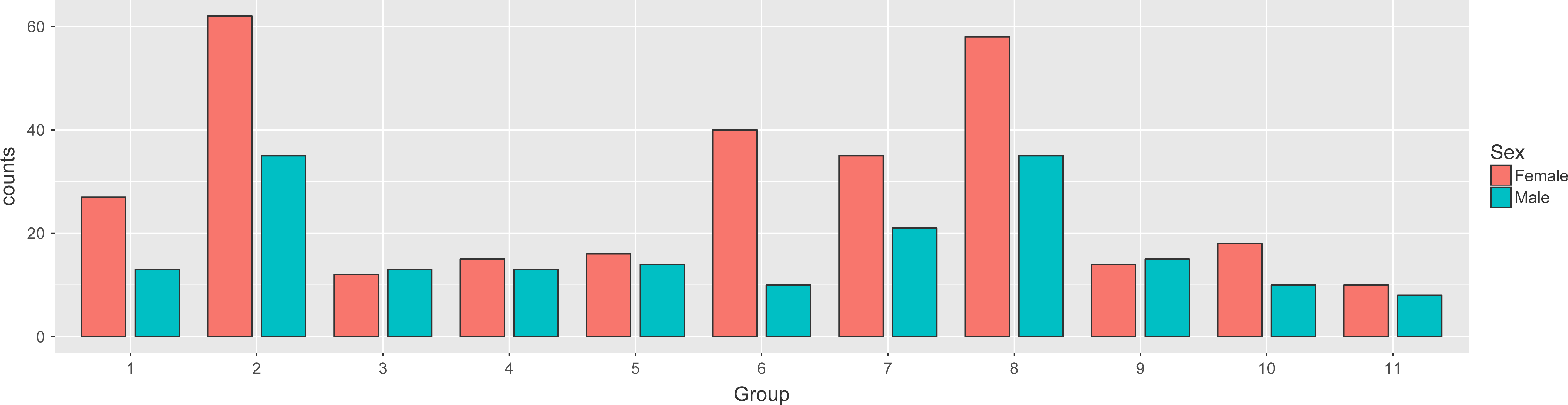
The distribution of participants based on age and gender. Participants were separated into 11 groups based on their age and displayed in male and female respectively. X-axis indicates the age group and Y-axis indicates the number of the participants. Blue bar indicates male participants and red bar indicates female participants, as well as the exact numbers of them are shown on the corresponding bar. Note that, for a relatively balance of distribution, the age span was set as 4 years in the first two groups, and 6 years in the rest of groups.

### Image Acquisitions

All of the data were collected at the Southwest University Center for Brain Imaging using a 3.0-T Siemens Trio MRI scanner (Siemens Medical, Erlangen, Germany). Each participant took part in 3D structural MRIand Resting-state fMRI scan; only for subgroups of participants have a task-fMRI scan (the task data is not part of this release). Additionally, to avoid the lasting effect of task fMRI on the resting fMRI, the resting state scan was performed before a particular task. For each participant, the3D structural MRI and resting state sequence were acquired in sequence during one session. The anatomical and resting state date was collected with the following parameters.

(1) **3D structural MRI** A magnetization-prepared rapid gradient echo (MPRAGE) sequence was used toacquire high-resolution Tl-weighted anatomical images (repetition time = 1,900 ms,echo time = 2.52 ms, inversion time = 900 ms, flip angle = 90 degrees, resolutionmatrix = 256 × 256, slices = 176, thickness =1.0 mm, voxel size = 1×1×1 mm^3^).
(2) **Resting-state fMRI** During the resting-state MRI scanning, the subjects were instructed to lie down,close their eyes, and rest without thinking about a specific thing, but refrain fromfalling asleep. The 8-min scan of 242 contiguous whole-brain resting-state functionalimages was obtained using gradient echo echo-planar-imaging (GRE-EPI) sequences with thefollowing parameters: slices = 32, repetition time (TR)/echo time (TE) = 2000/30 ms,flip angle = 90, field of view (FOV) = 220 × 220 mm, and thickness/slice gap = 3/1mm, and voxel size = 3.4 × 3.4 × 4 mm^3^.

### Code availability

We shared the code we used in the quality assessment (QA), voxel-based volume and functional connectivity analysis, and is freely available on github (https://github.com/Zhuang2KX/SALD).

### Date Records

This dataset is publicly available at the International Data-sharing Initiative (INDI) (All of MRI data can be accessed at http://fcon_1000.projects.nitrc.org/indi/retro/sald.html). Weremoved the facial information of each participant the S-MRI data (https://github.com/poldracklab/pydeface) and theNeuroimaging Informatics Technology Initiative (NIFTI) headers according toFCP/INDI policies. The contents and data structures of these packages are detailed asfollows:

### MRI data and demographic information

sMRI and rfMRI scans

Location: participant_id/scan_id/file.nii.gz

All the imaging data are organized according to BIDS criteria^9^. For more detailed information, please visit the following website: http://bids.neuroimaging.io/.

Demographic information

File format: ‘.csv’files

Basic demographic information including age, sex and handedness is provided in the ‘.csv’ file. Besides, the quality assessment measures to different scans were also included in other files.

### Quality Control Report

The folder quality-assessment-protocol compackage of quality assessment (QA) analysis resultsperformed in the present study for the structural and functional images. It contains csv files (namedqap_anatomical_spatial.csv,qap_functional_spatia.csv, and qap_functional_temporal.csv,respectively). Those files were generated by the Preprocessed Connectomes Project (PCP) Quality AssessmentProtocol and we didn’t change any part of the pipline. For more details about its procedure and the measures included, see the website of PCP Quality Assessment Protocol (http://preprocessed-connectomes-project.org/quality-assessment-protocol/). All data were made available to users regardless of data quality because there are no consensus criteria to determine what kind of MRI images should be excluded.

### Technical Validation

#### Results of QA measures

To quantitatively assess the quality of the MRI data, a series of widely used QA measures have been calculated. All measures computed by the PCP Quality Assessment Protocol can be found together with the data. Figures 2 and 3 indicate the distributions of the several representative QA measures of the structural MRI and resting-state fMRI, respectively, across participants. For more information about the QA measures, see uploaded csv files.

**Figure 2.**
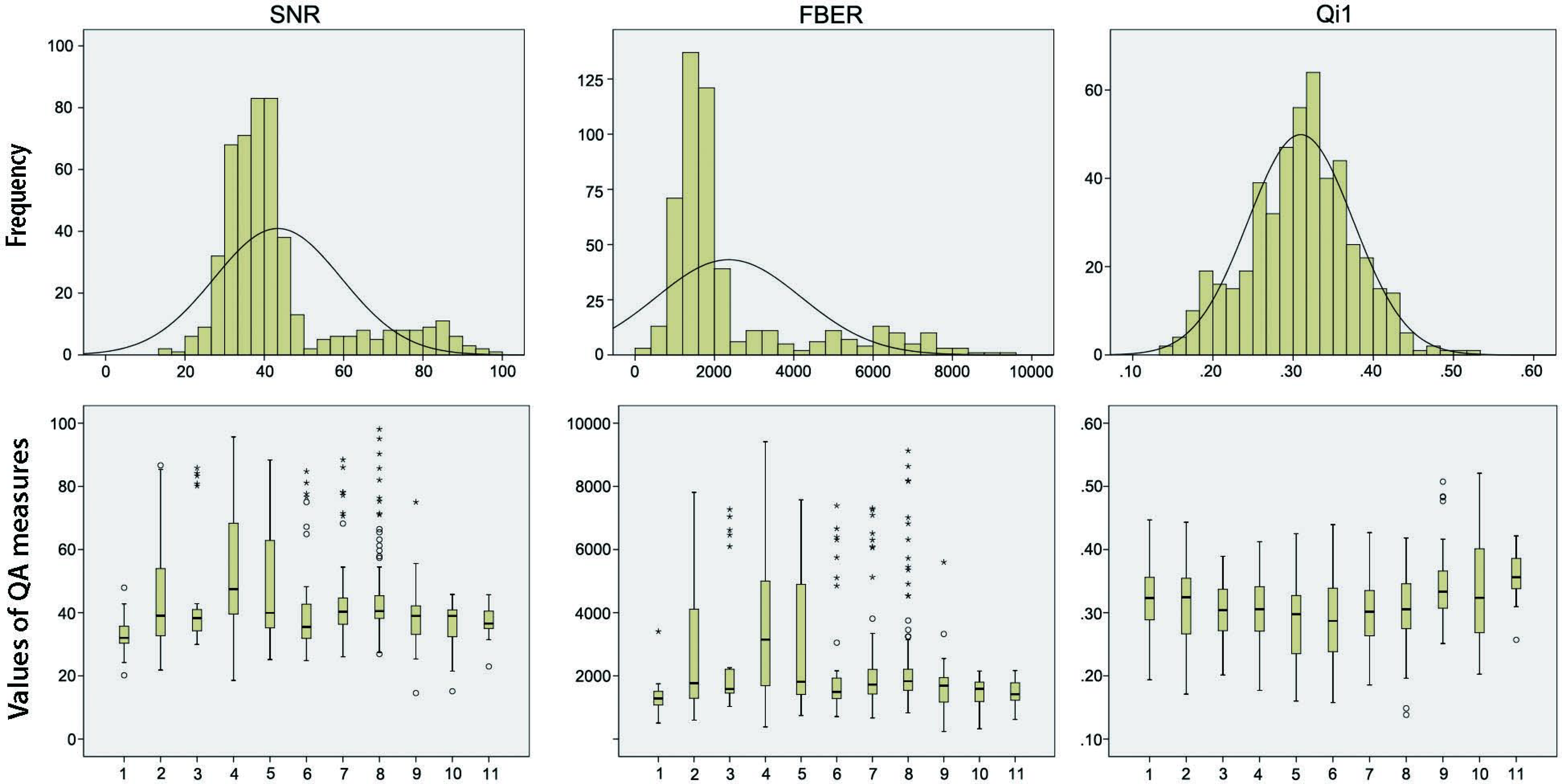
2 The distributions of the several representative QA measures of the structural MRI across all participants. **SNR** is the abbreviation of Signal-to-Noise Ratio. It indicates the mean intensity within gray matter divided by the standard deviation of the values outside the brain. Higher values are better (Magnotta, Friedman & BIRN, 2006). **FBER** is the abbreviation of Foreground to Background Energy Ratio. It indicates the variance of voxels inside the brain divided by the variance of voxels outside the brain. Higher values are better. **Qi1** means Percent Artifact Voxels, which implies proportion of voxels outside the brain with artifacts to the total number of voxels outside the brain. Lower values are better (Mortamet et al., 2009).

**Figure 3.**
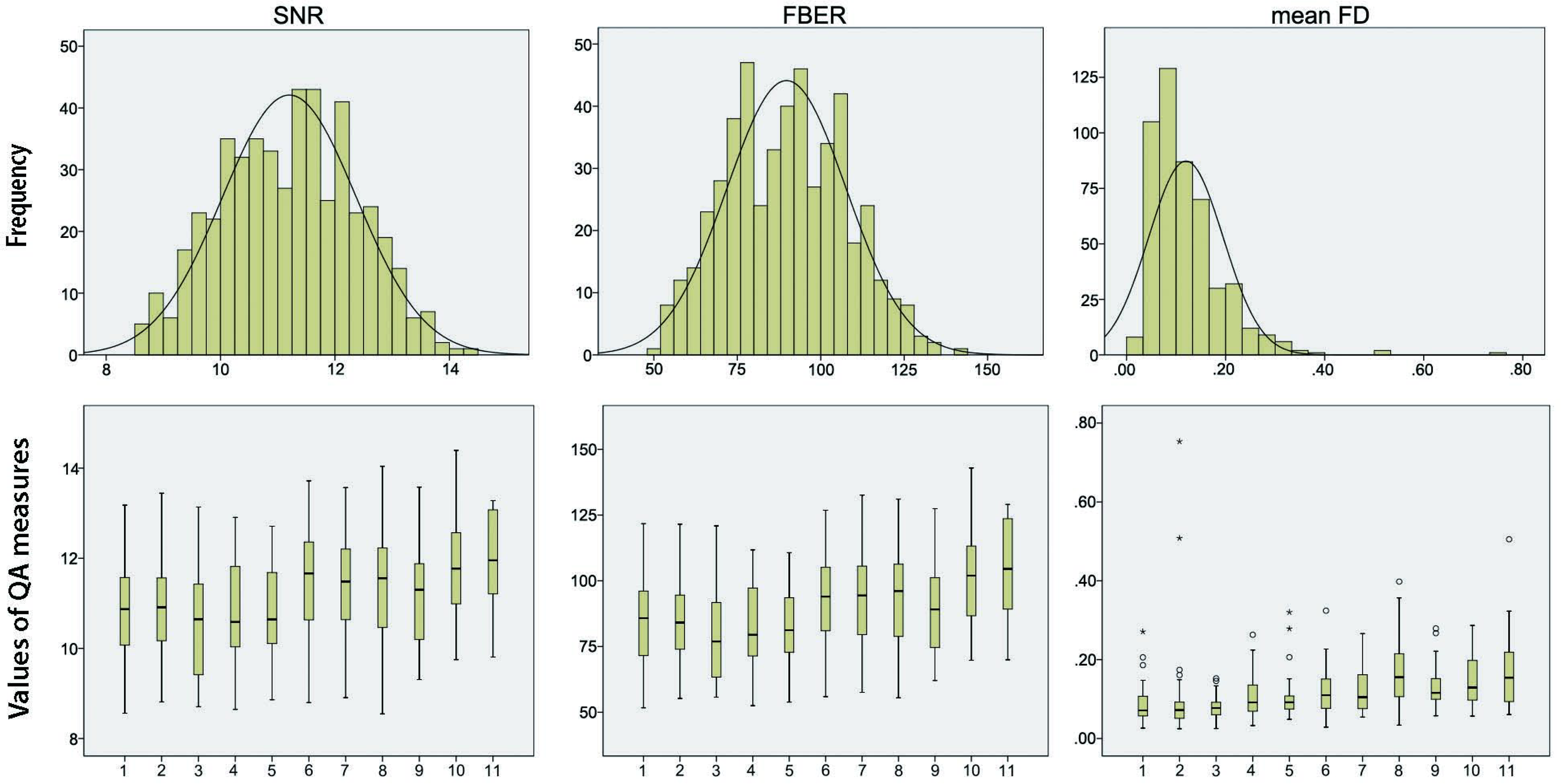
The distributions of the several representative QA measures of the resting-state fMRI across participants. **SNR** is the abbreviation of Signal-to-Noise Ratio. It indicates the mean intensity within gray matter divided by the standard deviation of the values outside the brain. Higher values are better □ Magnotta, Friedman & BIRN, 2006). **FBER** is the abbreviation of Foreground to Background Energy Ratio. It indicates the variance of voxels inside the brain divided by the variance of voxels outside the brain. Higher values are better. **Mean FD** means Mean Fractional Displacement-Jenkinson. It is a measure of subject head motion, which compares the motion between the current and previous volumes. This is calculated by summing the absolute value of displacement changes in the x, y and z directions and rotational changes about those three axes. The rotational changes are given distance values based on the changes across the surface of a 80mm radius sphere. Lower values are better (Jenkinson et al., 2002; Yan et al., 2013).

#### Relationship between age, head motion and signal-to-noise ratio (SNR)

To investigate the impact of head motion during the resting-state fMRI scanning on the overall quality of images and its association with age, we correlated the head motion (as measured by mean FD) with age and the SNR in the entire sample (N = 494). Results revealed no significant result was found in the relationship between mean FD and SNR (*r* = 0.055, *p* = 0.222). There is a significant and positive correlations exist between mean FD and age (*r* = 0.372, *p*< 0.001), and this relationship enhanced (*r* = 0.455, *p*< 0.001) after we removed 16 subjects who is the outliers of mean FD values. Figure 4 indicates these two correlations. The results suggested that head motion may increase with age and the head motion in this dataset didn’t significantly affect the overall quality of images in a linear trend.

**Figure 4.**
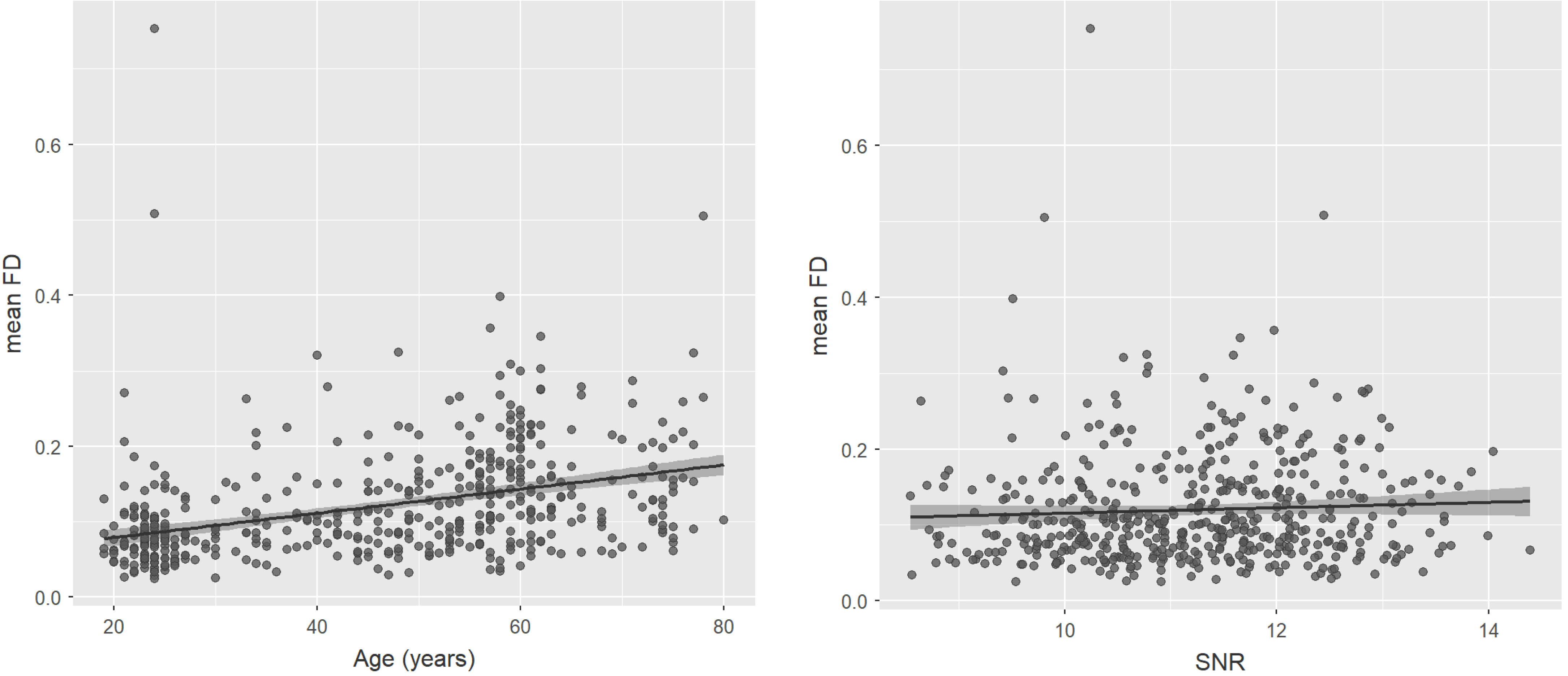
The affect of mean FD to age and SNR. The X-axes indicate age and SNR value respectively. The Y-axes indicate mean FD values.

#### Replication of previous findings

To test whether this dataset is technically valid,we tried to use the current data to replicate some previous findings. Here, sMRI and resting-state fMRI data were respectively analyzed based on this objective.

#### 3D structural MRI data

A large number of studies have reported that structural development during normal aging is accompanied with a declining trajectory of the total gray matter volume^10-14^; as well as, cortical gray matter volume was found to be a decline over adulthood^15-17^. Besides, the decreases were always reported to be most pronounced in the frontal and parietal lobes^6,12,15,18,19^. Here, we were attempting to replicate these robust findings in the current dataset.

The sMRI (1 × 1 × 1 mm^3^) data was preprocessed by using SPM8 (Welcome Department of Cognitive Neurology, Lodon, UK; www.fil.ion.ucl.ac.uk/spm). For better registration, all T1-weighted structural images were automatically co-registered to the anterior commissure-posterior commissure (AC-PC) by SPM8 based script. Then, a spatially adaptive nonlocal means (SANLM) denoising filter^20^ was used by VBM8 toolbox (http://www.neuro.uni-jena.de/vbm/download/). Next, using the unified segmentation procedure, the coregistered images from each participant were segmented into grey matter (GM), white matter and cerebrospinal fluid^21^. The GM images of each participant were spatially normalized to a study-specific T1-weighted template using a diffeomorphic nonlinear registration algorithm (DARTEL; diffeomorphic anatomical registration through exponentiated lie algebra).

The DARTEL registration involves: first computing the specific template based on the average tissue probability maps from all the participants; second warping each participant’s segmented maps into a specific template. In order to improve the alignment and achieve a more accurate inter-subject registration, the procedure was repetitively conducted until a best study-specific template was generated. Subsequently, registered images were transformed to Montreal Neurological Institute (MNI) space and a further modulation was conducted to preserve the volume of GM. Finally, a 6-mm full width at half-maximum (FWHM) Gaussian kernel was applied to smooth the modulated GM images.

We first used Pearson correlation to detect the relationship between age and total gray matter volume (GMV). Then, multiple linear regressions were used to determine GMV regions that were associated with age, controlling for total GMV. To avoid edge effects around the borders between GM and WM, we used explicit masking to restrict the search volume. The explicit masking was achieved by the SPM Masking Toolbox (http://www.cs.ucl.ac.uk/staff/g.ridgway/masking/). This approach reduced the risk of false negatives caused by overly restrictive masking, as potentially interesting voxels may be excluded from the statistical analysis^22^. For the regression analysis, we used the family-wise error (FWE) of *p*< 0.05 at the whole brain level and ≥ 20 contiguous voxels as a threshold to correct for multiple comparisons.

The results indicated that age is significantly correlated with total GMV (*r* = ‐0.305, *p*< 0.001). Almost all areas of the cerebral cortex exhibited a significant age-related decline in GMV. In addition, frontal, parietal and temporal lobes showed most pronounced function, which to a large extent confirmed previous findings (Fig. 5). However, in accordance with one prior research^6^, we found that occipital regions were less affected by age.

**Figure 5.**
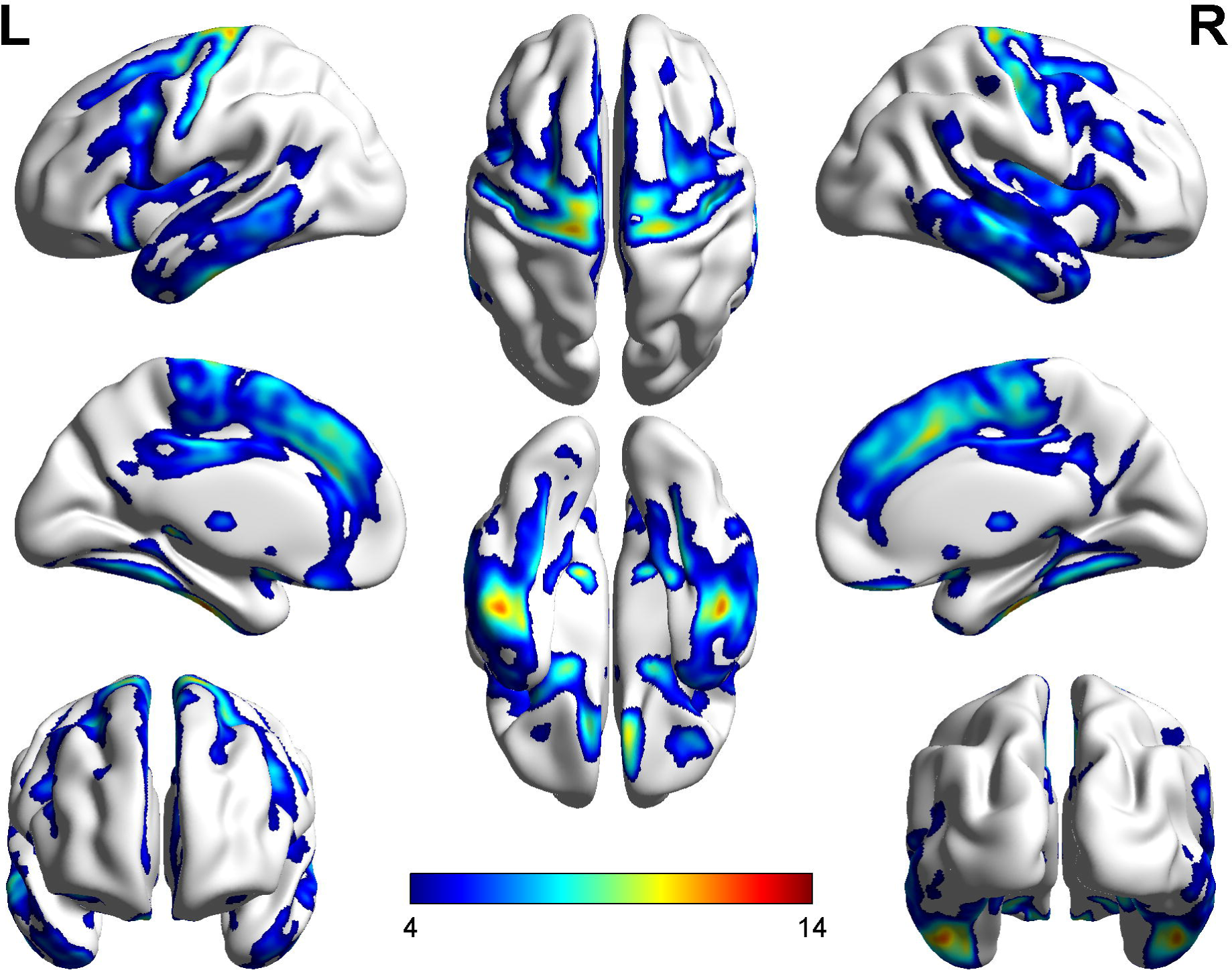
Brain regions with GMV reduction in normal aging. L-R means from left brain hemisphere to right hemisphere.

#### Resting-state fMRI data

There is a widely reported finding indicated that clear segmentation between neural systems would lose consistently over the course of normal human aging: many intrinsic functional connectivity brain networks gradually become less internally coherent with age^7,23-25^. For the attempt to replicate this finding, the current dataset was used to describe the changing trajectories of within-system connectivity along with age.

The resting-state fMRI data were preprocessed using Data Processing Assistant for Resting-State fMRI (DPARSF_V4.2, http://resting-fmri.sourceforge.net/) implemented in the MATLAB 2014a (Math Works, Natick, MA, USA) platform. The first 10 volumes of the functional images were discarded to account for signal equilibrium and the participants^1^ adaptation to their immediate environment. The remaining 232 scans were corrected for slice timing, and then realigned to the middle volume to correct for head motion. Participant with head motion exceeding 2.0 mm in any dimension throughout the course of scans was discarded from further analysis. Subsequently, registered images were spatially normalized to Montreal Neurological Institute (MNI) template (resampling voxel size = 3 × 3 × 4 mm^3^). Next, nuisance signals representing motion parameters, global signals, white matter, and cerebrospinal fluid signals were regressed out in order to control the potential impact of physiological artifacts. Here, we used the Friston 24-parameter model, including 6 motion parameters, 6 temporal derivatives, and their squares^26,27^ to regress out head motion effects. This approach is based on recent research demonstrating that higher-order models are more effective at reducing the effects of head movements^28-30^. Then, after the spatial smoothing (full width at half maximum = 6 mm Gaussian kernel), bandpass filtering (0.009–0.08 Hz) was performed. These preprocessing steps were followed by the standard protocol published^29^.

Whole-brain functional connectome were constructed for each subject as a 264 × 264-node graph, labeled by functional systems^28^. Edge weights were calculated as the Fisher z-transformed correlation (Pearson’s r) between each pair of nodes, and negatively weighted edges were removed from each correlation matrix to eliminate potential misinterpretation of negative edge weights. For a specific system, within-system connectivity was calculated as the mean node-to-node z-value of all nodes of that system to each other. The mean within-system connectivity means the average value of within-system connectivity over all of the systems.

The results indicated that mean within-system connectivity would decrease with age. When we applied linear and nonlinear (second-degree polynomial) to within-system connectivity, we found that the age function was fit significantly both by the linear model (adjusted *R*^*2*^= 0.177, *p*< 0.001) and nonlinear model (adjusted *R*^*2*^= 0.195, *p*< 0.001). While, the quadratic model had a higher *R*^*2*^ than the linear model, which implied a preservation of within-system connectivity during the early adult lifespan (Fig. 6).

**Figure 6.**
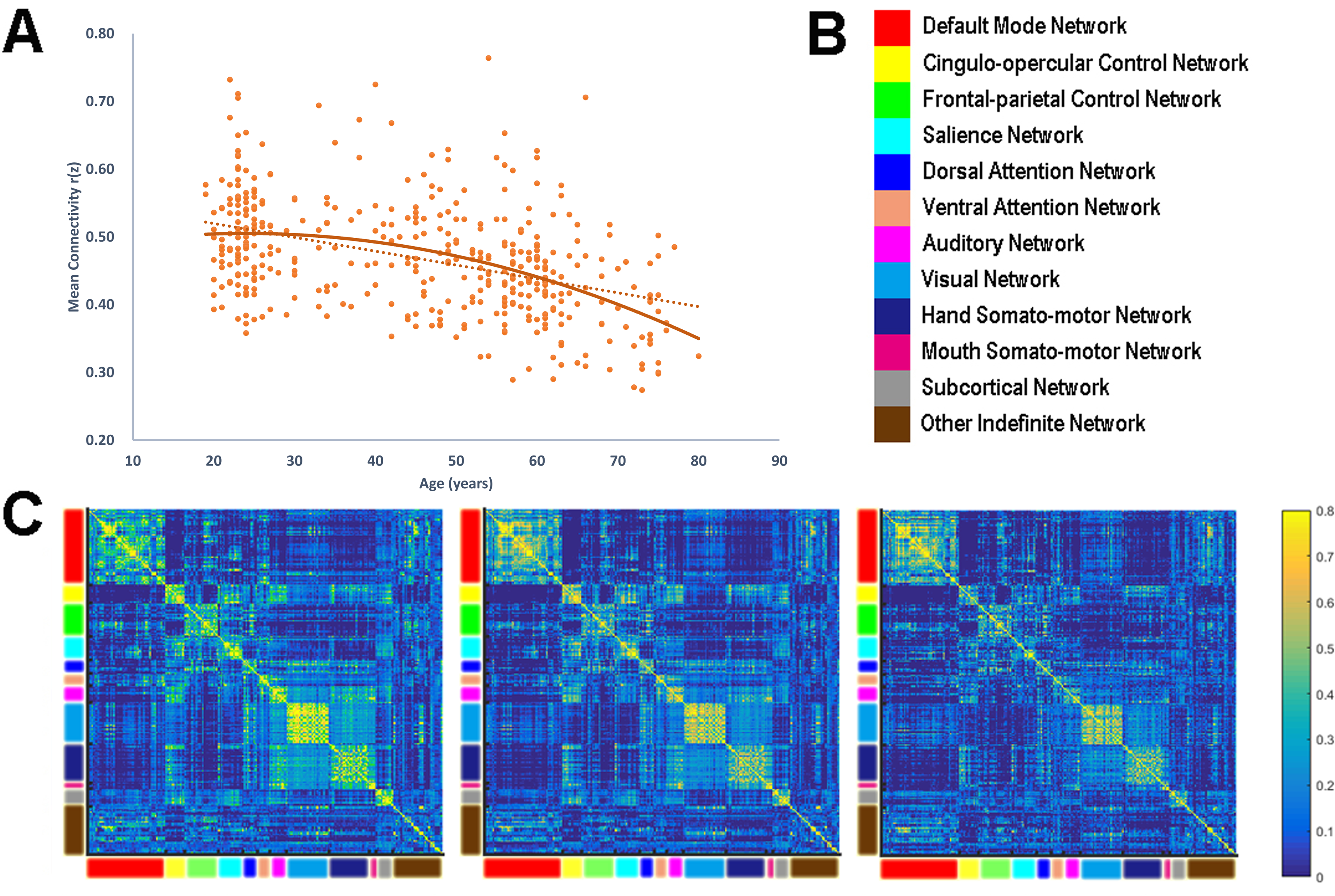
Within-system connectivity decline with aging. **A** demonstrates the negative correlation between age and mean connectivity. **B** displays the different brain networks (Power et al., 2011) involved in this analysis. The mean connectivity in A was calculated by averaging the intrinsic functional connectivity within each of the networks. **A** displays the functional connectivity matrices of three representative age groups. The networks were arranged as the same order as **B**. It can be seen that the within-system connectivity apparently declines with aging.

## Usage Notes

We encourage other labs to use this dataset in publication under the requirement of citing the present data descriptor. All data is free to download from the International Data-sharing Initiative (http://fcon_1000.projects.nitrc.org/indi/retro/sald.html). The results of quality analysis measure are available for free download and use according to consensus criteria to determine what kind of MRI images should be excluded.We hope that all users of the data will acknowledge the original authors by citing this publication.

## Acknowledgements

This dataset was supported by:The National Natural Science Foundation of China (31571137; 31500885), National Outstanding young people plan, the Program for the Top Young Talents by Chongqing, the Fundamental Research Funds for the Central Universities (SWU1509383; SWU1509451; SWU1609177), Natural Science Foundation of Chongqing (cstc2015jcyjA10106),Fok Ying Tung Education Foundation (151023) to J.Q, D.T.W and W.J.Y. We are grateful to all of graduate students who contributed their time and wisdom to this data repository, including but not limited to Xin Hou, Junyi Yang, Xue Tian, Hui Huang, Xiayun Man, and Yadan Li. We are also thanking Michael Milham, David O'Connor, Lei Ai, and Bonhwang Koo for the data repository.

## Author Contributions

Conception and design: J.Q.; Analysis and interpretation: D.T.W, K.X.Z, Q.L.C,and J.Q.; Data collection: Q.L.C, W.J.Y, W. L, K.C.W, and J.Z.S. Writing the article: D.T.W, K.X.Z and J.Q.

## Data Citations

1. Wei, D.-T.,Zhuang, K.-X., Chen, Q.-L.,Liu, W.,Qiu, J.International Data-sharing lnitiative http://fcon_1000.projects,nitrc.org/indi/retro/sald.html (2017).

## References

1 Krogsrud, S. K. et al. Changes in white matter microstructure in the developing brain—A longitudinal diffusion tensor imaging study of children from 4 to llyears of age. NeuroImage 124, 473-486 (2016).

2 Maniega, S. M. et al. White matter hyperintensities and normal-appearing white matter integrity in the aging brain. Neurobiology ofaging 36,909-918 (2015).

3 Hazlett, H. C. et al. Early brain development in infants at high risk for autism spectrum disorder. Nature 542, 348-351 (2017).

4 Dosenbach, N. U. et al. Prediction of individual brain maturity using fMRI. Science 329, 1358-1361 (2010).

5 Finn, E. S. et al. Functional connectome fingerprinting: identifying individuals using patterns of brain connectivity. Nature neuroscience 18, 1664-1671 (2015).

6 Ziegler, G. et al. Brain structural trajectories over the adult lifespan. Human brain mapping 33, 2377-2389 (2012).

7 Chan, Μ. Y., Park, D. C., Savalia, N. Κ., Petersen, S. E. & Wig, G. S. Decreased segregation of brain systems across the healthy adult lifespan. Proceedings of the National Academy of Sciences 111, E4997-E5006 (2014).

8 Taylor, J. R. et al. The Cambridge Centre for Ageing and Neuroscience (Cam-CAN) data repository: structural and functional MRI, MEG, and cognitive data from a cross-sectional adult lifespan sample. Neuroimage (2015).

9 Gorgolewski, K. J. et al. The brain imaging data structure, a format for organizing and describing outputs of neuroimaging experiments. Scientific Data 3, 160044 (2016).

10 Allen, J. S., Bruss, J., Brown, C. K. & Damasio, H. Normal neuroanatomical variation due to age: the major lobes and a parcellation of the temporal region. Neurobiology of aging 26, 1245-1260 (2005).

11 Giorgio, A. et al. Age-related changes in grey and white matter structure throughout adulthood. NeuroImage 51, 943-951 (2010).

12 Good, C. D. et al. in Biomedical Imaging, 2002. 5th IEEE EMBS International Summer School on. 16 pp. (IEEE).

13 Grieve, S. M., Clark, C. R., Williams, L. M., Peduto, A. J. & Gordon, E. Preservation of limbic and paralimbic structures in aging. Human brain mapping 25, 391-401 (2005).

14 Hasan, Κ. M. et al. Development and organization of the human brain tissue compartments across the lifespan using diffusion tensor imaging. Neuroreport 18, 1735-1739 (2007).

15 Kalpouzos, G. et al. Voxel-based mapping of brain gray matter volume and glucose metabolism profiles in normal aging. Neurobiology of aging 30, 112-124 (2009).

16 Sullivan, E. V., Rosenbloom, M., Serventi, K. L. & Pfefferbaum, A. Effects of age and sex on volumes of the thalamus, pons, and cortex. Neurobiology of aging 25, 185-192 (2004).

17 Walhovd, Κ. B. et al. Effects of age on volumes of cortex, white matter and subcortical structures. Neurobiology of aging 26, 1261-1270 (2005).

18 Resnick, S. M., Pham, D. L., Kraut, M. A., Zonderman, A. B. & Davatzikos, C. Longitudinal magnetic resonance imaging studies of older adults: a shrinking brain. Journal of Neuroscience 23, 3295-3301 (2003).

19 Smith, C. D., Chebrolu, H., Wekstein, D. R., Schmitt, F. A. & Markesbery, W. R. Age and gender effects on human brain anatomy: a voxel-based morphometric study in healthy elderly. Neurobiology of aging 28,1075-1087 (2007).

20 Manjón, J. V., Coupé, P., Martí-Bonmatí, L.., Collins, D. L. & Robles, M. Adaptive non-local means denoising of MR images with spatially varying noise levels. Journal of Magnetic Resonance Imaging 31, 192-203 (2010).

21 Ashburner, J. & Friston, K. J. Unified segmentation. Neurolmage 26,839-851 (2005).

22 Ridgway, G. R. et al. Issues with threshold masking in voxel-based morphometry of atrophied brains. NeuroImage 44, 99-111 (2009).

23 Contreras, J. A., Goñi, J., Risacher, S. L., Sporns, O. & Saykin, A. J. The structural and functional connectome and prediction of risk for cognitive impairment in older adults. Current behavioral neuroscience reports 2, 234-245 (2015).

24 Dennis, E. L. & Thompson, P. M. Functional brain connectivity using fMRI in aging and Alzheimer’s disease. Neuropsychology review 24,49-62 (2014).

25 Sala-Llonch, R.., Bartrés-Faz, D. & Junqué, C. Reorganization of brain networks in aging: a review of functional connectivity studies. Frontiers in psychology 6 (2015).

26 Friston, K. J., Williams, S., Howard, R., Frackowiak, R. S. & Turner, R. Movement-related effects in fMRI time-series. Magnetic resonance in medicine 35,346-355 (1996).

27 Satterthwaite, T. D. et al. An improved framework for confound regression and filtering for control of motion artifact in the preprocessing of resting-state functional connectivity data. NeuroImage 64, 240-256 (2013).

28 Power, J. D., Barnes, K. A., Snyder, A. Z., Schlaggar, B. L. & Petersen, S. E. Spurious but systematic correlations in functional connectivity MRI networks arise from subject motion. NeuroImage 59, 2142-2154 (2012).

29 Yan, C.-G., Wang, X.-D., Zuo, X.-N. & Zang, Y.-F. DPABI: data processing & analysis for (resting-state) brain imaging. NeuroinformaticslQ, 339-351 (2016).

30 Yan, C.-G. et al. A comprehensive assessment of regional variation in the impact of head micromovements on functional connectomics. Neurolmage 76,183-201 (2013).

